# Evolution and Human Neural Individuality

**DOI:** 10.64898/2026.08.26.747255

**Authors:** Noga Yair, Yael Coldham, Ido Tavor, Yair Bar-Haim

**Affiliations:** Sagol School of Neuroscience, Tel-Aviv University, Tel Aviv, Israel; School of Psychological Sciences, Tel-Aviv University, Tel Aviv, Israel; Gray School of Medical Sciences, Gray Faculty of Medical & Health Sciences, Tel Aviv University, Tel Aviv, Israel

## Abstract

Individuality is a defining feature of human biology. The functional network architecture of the human brain harbors person-specific qualities and forms individualized connectivity profiles that function as a neural ‘fingerprint’, both stable and unique across time. Here, using fMRI data from 431 Human Connectome Project participants, we examined whether neural individuality is more strongly exhibited in brain regions bearing signatures of recent human evolution. We calculated region-wise fingerprinting accuracy and associated it with four properties of evolutionary cortical organization: cortical expansion, myelin content estimate (T1w/T2w), human-specific gene-expression profiles, and functional homology to other primates. Across all four measures, neural individuality was strongest in cortical areas showing greater evolutionary novelty in humans, particularly frontoparietal control and default mode networks, and weaker in more conserved primary regions. Our findings connect evolutionary variation across species with stable functional variation among individuals.

## Main text

The coherent sense of self is thought to depend on the brain’s capacity to integrate cognition, perception, emotion, and self-representation across time. Whole-brain functional connectivity provides a systems-level substrate for this integration^1,2^. The large-scale networks supporting cognitive, affective, and self-referential processes exhibit highly individualized connectivity profiles that reliably distinguish individuals across time and context^3,4^. Using Human Connectome Project data (HCP)^5^, Finn et al.^3^ showed that connectivity profiles derived from functional magnetic resonance imaging (fMRI) constitute stable neural ‘fingerprints’, accurately identifying individuals across sessions separated by two days. Subsequent studies have demonstrated that these ‘fingerprints’ persist across resting-state and task conditions^3,4^ and are associated with individual differences in cognitive abilities and psychopathology^6,7^. The functional connectome therefore captures a fundamental dimension of human neural individuality. However, the evolutionary origins of this neural individuality remain unknown.

Connectome fingerprinting is supported disproportionately by frontoparietal regions^3,8^. These regions exhibit a pronounced role in higher-order cognition and self-referential processing, capacities central to human cognition^1,9^. Unlike primary sensory and motor regions, whose organization is tightly constrained by shared perceptual and motor demands, frontoparietal association cortices may permit greater interindividual variability. Consistent with this view, individual-specific variation in functional network organization is particularly pronounced in these regions^10^ which support higher-order functions including cognitive control, working memory, and decision-making^11,12^. Comparative neuroanatomical evidence further indicates that association cortices, particularly prefrontal and parietal regions, underwent substantial expansion and reorganization during human evolution^13,14^. For example, gyrification is especially pronounced in association cortices, with the largest anatomical differences between humans and nonhuman primates observed in prefrontal regions^15^. Thus, the regions that most reliably distinguish individuals through functional connectivity may be precisely those in which human evolution created the greatest capacity for flexible, individualized network organization.

We hypothesized that the neural basis of human individuality, as indexed by functional connectome fingerprinting, is driven most strongly by phylogenetically recent cortical systems shaped by human-specific evolutionary adaptations. To test this hypothesis, we compared multimodal indices of evolutionary novelty across humans, rhesus macaques (*Macaca mulatta*), and chimpanzees (*Pan troglodytes*), whose lineage diverged from that of humans approximately 25 and 6 million years ago (MYA), respectively^16^ (Figure 1A). We used a four-pronged approach linking connectivity-based fingerprinting to regional variation in: (1) cortical expansion relative to other primates; (2) estimated myelin content; (3) human-specific gene-expression profiles; and (4) functional homology across primate species. We further derived a Composite Evolutionary Index (CEI) integrating these measures into a single region-level estimate of evolutionary-recent cortical organization. If regions supporting individual identification systematically overlap with regions that underwent the greatest evolutionary specialization in humans, this would suggest that the emergence of uniquely human cortical organization also increased the brain’s capacity for individualized functional architecture.

**Figure 1.**
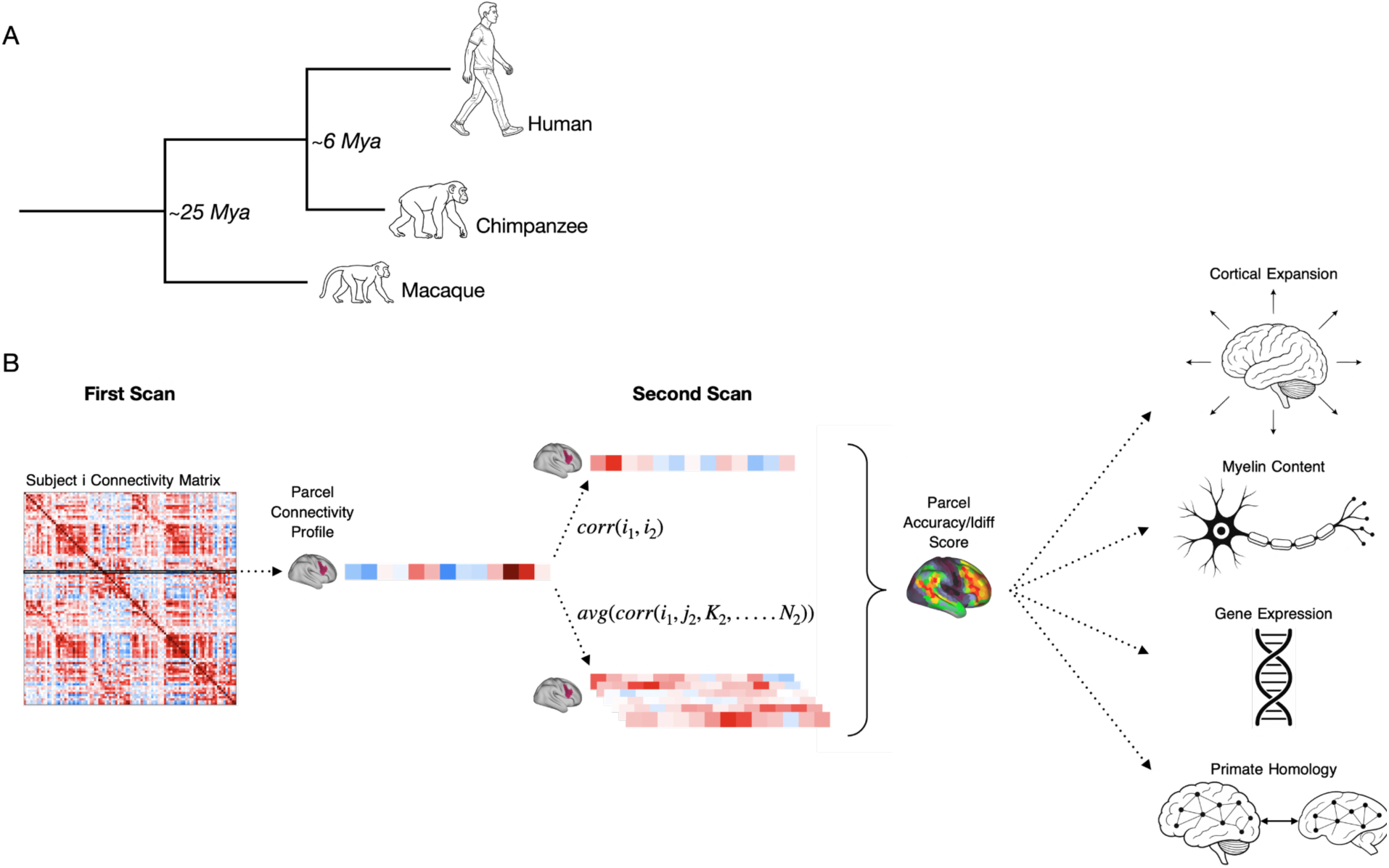
Conceptual overview of the evolutionary fingerprinting framework. (A) Phylogenetic tree illustrating the evolutionary divergence of humans, chimpanzees, and macaques. (B) Schematic overview of the analytical pipeline. Resting-state functional connectivity matrices from individual participants were used to calculate parcel-wise connectome fingerprinting metrics, including identification accuracy and differential identifiability (Idiff). For each cortical parcel, evolutionary and neurobiological properties were quantified using chimpanzee-to-human cortical expansion, cortical myelination estimate (T1w/T2w), HAR-BRAIN gene expression, and the Functional Connectivity Homology Index (FCHI). Associations between parcel-wise fingerprinting metrics and each evolutionary measure were then evaluated.

## Neural Fingerprinting

We analyzed resting-state functional MRI (rsfMRI) data from 431 unrelated participants in the HCP S1200 release^5^. Each participant completed two rsfMRI sessions on separate days. The cortex was parcellated into 1,000 regions using the Schaefer atlas^17^, with each parcel assigned to one of seven canonical functional networks^18^. For each participant and session, we extracted the mean rsfMRI time series for each parcel and correlated it with the time series of all other parcels, generating a parcel-specific functional connectivity profile (Figure 1B).

Rather than using the whole connectome for identification, we quantified the fingerprinting capacity of each cortical parcel based on its specific connectivity profile (Figure 1B). For each participant, the connectivity profile from one session was correlated with the corresponding parcel-specific profiles of all participants from the other session. Fingerprinting success was recorded when the highest cross-session correlation was within the same participant. Parcel-wise fingerprinting accuracy was then calculated as the proportion of successful identification across participants. Higher accuracy indicated that a parcel’s connectivity profile was both stable within individuals and sufficiently distinctive to reliably identify them across sessions.

As a complementary measure, we calculated differential identifiability^4^ (Idiff), defined as the difference between each participant’s cross-session self-correlation and their mean correlation with all other participants. Higher Idiff values indicate that a parcel’s connectivity profile more strongly distinguishes an individual from others, irrespective of whether exact fingerprinting identification was achieved. Together, fingerprinting accuracy and Idiff provide complementary measures of parcel-level individuality. Accuracy quantifies discrete identification success. Idiff quantifies the continuous separation between within- and between-participant similarity. Parcel-level accuracy is shown in Figures 2-4, and parcel-wise Idiff in Supplementary Figure S1.

**Figure 2.**
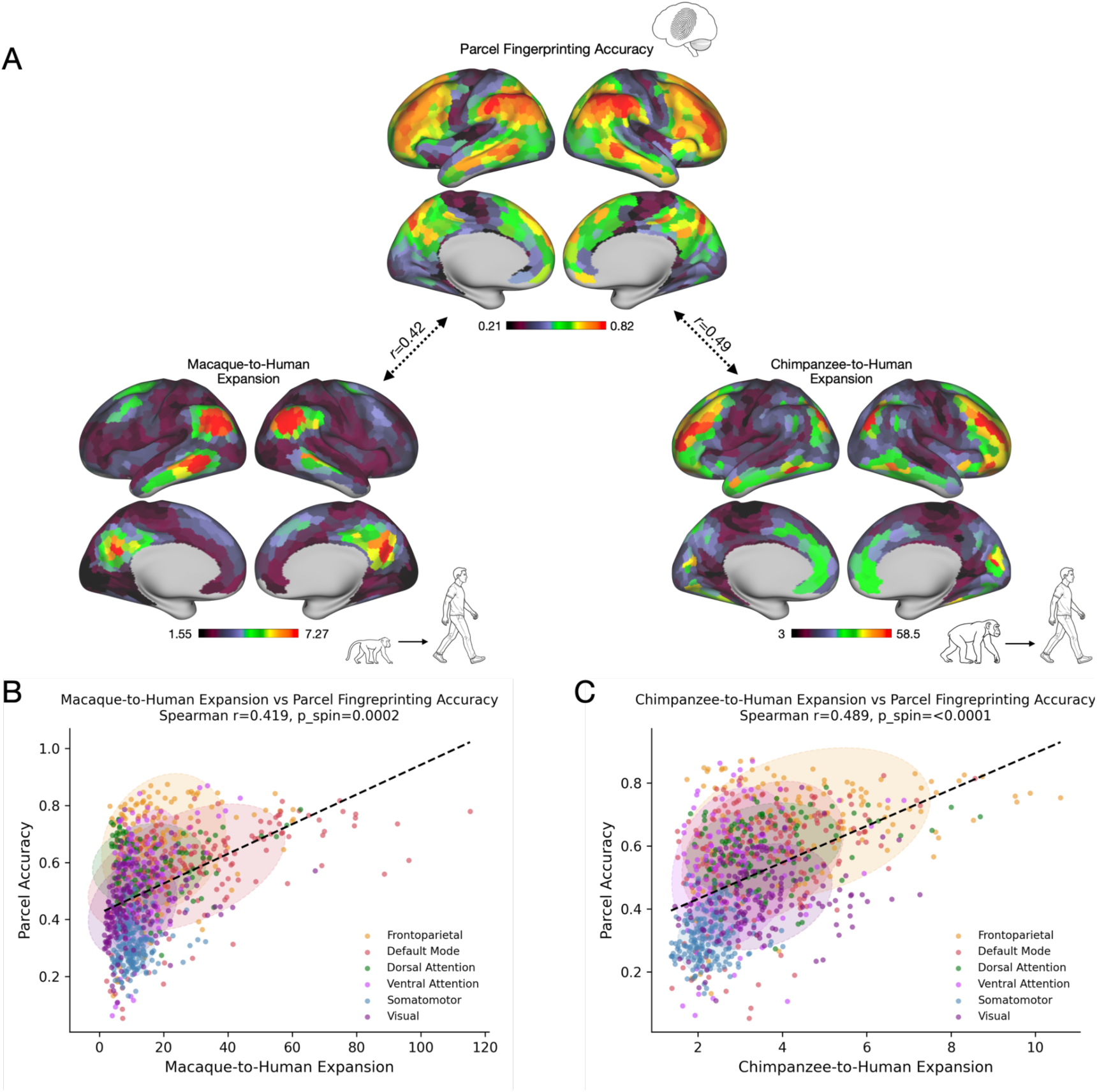
Cortical expansion is associated with parcel-level fingerprinting accuracy. (A) Parcel-wise fingerprinting accuracy (top) compared to macaque-to-human (left) and chimpanzee-to-human (right) cortical expansion, mapped onto the Schaefer 1,000-parcel cortical surface. Warmer colors indicate higher values. (B) Association between macaque-to-human cortical expansion and parcel-level fingerprinting accuracy. (C) Association between chimpanzee-to-human cortical expansion and parcel-level fingerprinting accuracy. Each point represents one cortical parcel and is colored according to its canonical functional network. Shaded ellipses indicate the distribution of parcels within each canonical network, and dashed lines show the overall linear fit for visualization. Fingerprinting accuracy was positively and significantly associated with both macaque-to-human expansion (*r* = 0.419, *p*_spin_ < 0.001) and chimpanzee-to-human expansion (*r* = 0.489, *p*_spin_ < 0.0001).

Analyses focused on the neocortex, which underwent substantial expansion and reorganization during human evolution and supports many high-order cognitive and affective functions^19,20^. We excluded parcels assigned to the Yeo limbic network due to their concentration in the bilateral temporal poles, ventral anterior temporal lobes, and orbitofrontal cortex^18^, regions that are particularly susceptible to signal loss and geometric distortion in conventional fMRI acquisition owing to their proximity to paranasal air cavities^21^. These regions were excluded because their systematically lower fMRI signal quality can reduce the reliability of functional connectivity estimates, potentially confounding comparisons of parcel-level fingerprinting performance.

## Cortical expansion

The disproportionate expansion of the cerebral cortex is a defining feature of human evolution. Cortical expansion has been linked to anatomical and functional specialization, with interconnected neural systems tending to evolve in concert^19^. Accordingly, regional expansion covaries with multiple cortical properties, including molecular architecture^22^, functional organization^23^, and microstructure^24^. However, cortical expansion has been highly heterogeneous across the cortex^25^, with frontal, temporal, and parietal association cortices expanding substantially more than relatively conserved primary sensory regions^14^. This evolutionary gradient raises the possibility that recently expanded cortical regions also possess greater capacity for individualized functional organization, making them disproportionately important for defining person-specific functional connectomes.

We examined whether regional variation in neocortex connectome fingerprinting was associated with cortical expansion during primate evolution. Macaque-to-human and chimpanzee-to-human cortical expansion maps were obtained from Xu et al.^23^ and Wei et al.^22^, respectively. Consistent with previous work^26^, both maps showed pronounced primate-to-human cortical expansion in higher-order association cortex, particularly in frontoparietal regions. However, the spatial distribution differed across species. Expansion relative to macaques was greater in parietal regions, whereas expansion relative to chimpanzees, humans’ closest living relatives, was concentrated in frontal regions (Figure 2A). This shift is consistent with the view that different association systems underwent peak expansion at different stages of primate evolution.

We hypothesized that cortical regions exhibiting greater evolutionary expansion in humans relative to other primates would contribute more strongly to neural fingerprinting than relatively conserved regions. To test this hypothesis, we averaged macaque-to-human and chimpanzee-to-human expansion values within each Schaefer parcel and correlated these estimates with parcel-wise fingerprinting accuracy using Spearman rank correlations. Statistical significance was assessed using 10,000 spatial spin permutations to account for spatial autocorrelation across the cortical surface.

Fingerprinting accuracy was positively associated with both macaque-to-human cortical expansion (Spearman’s *r* = 0.419, *p*_spin_ < 0.001; Fig. 2B) and chimpanzee-to-human cortical expansion (*r* = 0.489, *p*_spin_ < 0.0001; Fig. 2B). Thus, cortical regions that underwent greater evolutionary expansion in human lineage exhibited more distinctive individual-specific functional connectivity profiles. Analyses using Idiff yielded convergent results (Supplementary Figure S1).

## Myelin content

Evolutionary differentiation of the human cortex is also reflected in regional patterns of cortical myelination^24^. Throughout this study, “myelin” refers to T1w/T2w ratio, a widely used MRI-derived proxy for cortical myelin content^27^. Comparative analyses have shown that, relative to chimpanzees and macaques, the human cortex exhibits disproportionate expansion of lightly myelinated cortices, whereas heavily myelinated regions are comparatively conserved^20,28^. We therefore hypothesized that lightly myelinated cortical regions would exhibit greater fingerprinting accuracy than more heavily myelinated regions.

For each cortical parcel, mean myelin (T1w/T2w) was estimated by averaging vertex-wise values from the HCP cortical myelin map within the Schaefer 1,000-parcel atlas (Fig. 3A). Parcel-wise myelin estimates were then correlated with fingerprinting accuracy using Spearman rank correlations, with statistical significance assessed using 10,000 spatial spin permutations to account for spatial autocorrelation across the cortical surface.

**Figure 3.**
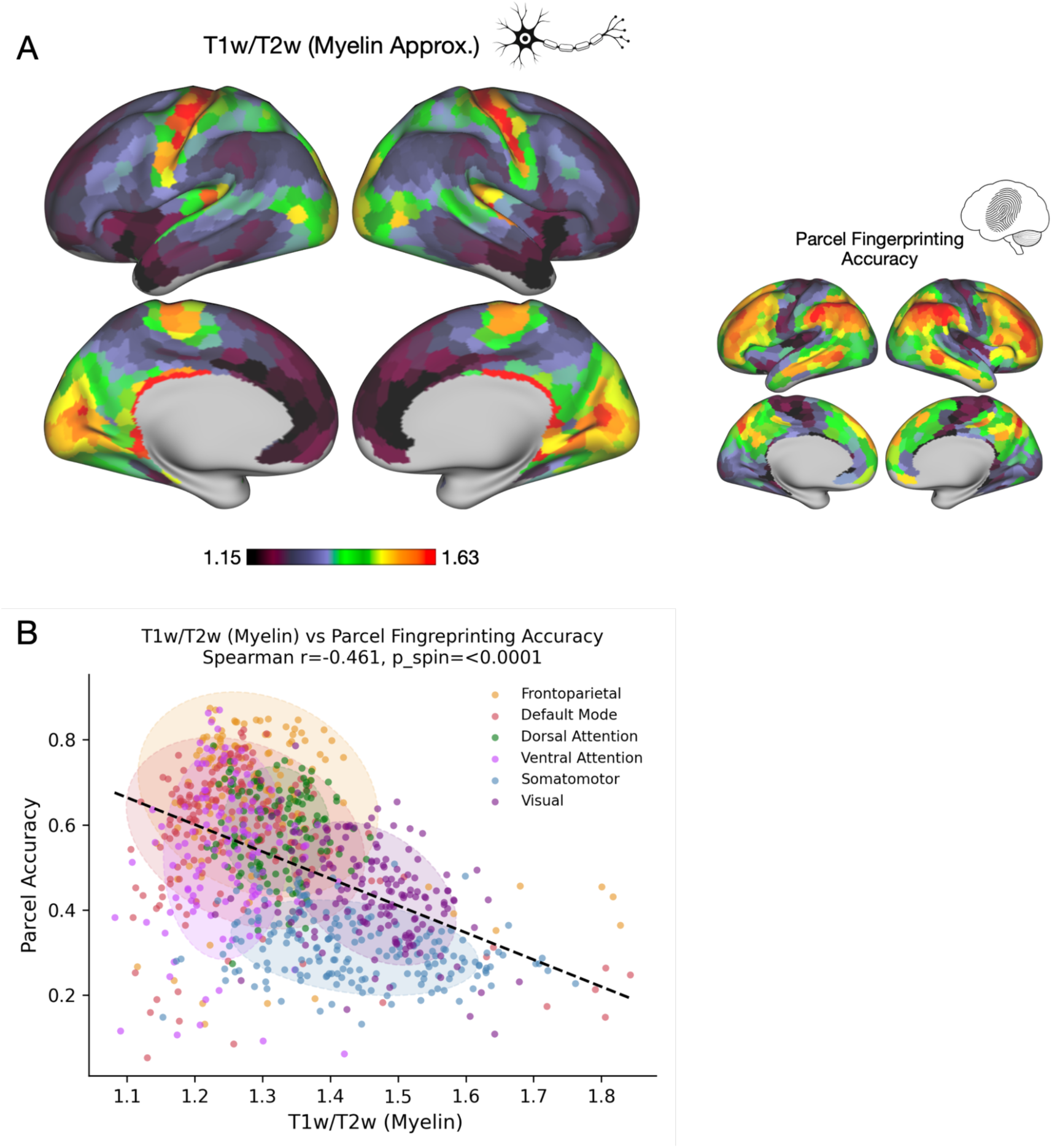
Cortical “myelin” (T1w/T2w) is inversely correlated with fingerprinting accuracy. (A) Spatial distribution of cortical myelin content mapped onto the Schaefer 1,000-parcel cortical surface. The fingerprinting-accuracy map, as also shown in Fig. 2A, is reproduced at a smaller scale (right) for visual comparison. Warmer colors indicate higher values. (B) Association between parcel-level myelin and fingerprinting accuracy. Each point represents one cortical parcel and is colored by canonical functional network. Shaded ellipses indicate the distribution of parcels within each network, and the dashed line shows the overall linear fit for visualization. Fingerprinting accuracy is negatively and significantly associated with cortical myelin (*r* = −0.461, *p*_spin_ < 0.0001), indicating that more lightly myelinated cortical regions exhibit more individually distinctive functional connectivity profiles.

As hypothesized, fingerprinting accuracy was negatively associated with cortical myelin content (*r* = -0.461, *p_spin_* < 0.0001; Fig. 3B), indicating that lightly myelinated cortical regions exhibited more distinct individual-specific functional connectivity profiles than heavily myelinated regions. Analyses using Idiff yielded convergent results (Supplementary Figure S1).

## Gene expression

Although humans share much of their genomic architecture with other mammals, species differences in gene regulation and expression are thought to underlie uniquely human features of brain development^20,28^ and adult cortical organization^29,30^. Distinct transcriptional profiles across the neocortex are thought to specify regional cortical identity during development^31^. We therefore asked whether cortical regions exhibiting greater human-specific gene-expression profiles also contribute more strongly to individual differences in functional network organization, as indexed by neural fingerprinting.

Human accelerated regions (HARs) are genomic loci that are highly conserved across vertebrates, but exhibit marked sequence divergence in humans, making them candidate regulatory elements underlying human-specific traits^32^. Although predominantly non-coding, HARs are enriched genes involved in neurogenesis, transcriptional regulation, and cortical development, suggesting that human brain evolution was driven in part by changes in gene regulation. Consistent with this view, Wei et al.^22^ reported that HAR-BRAIN genes are preferentially expressed in frontoparietal association cortices in humans, implicating these genes in the evolution of higher-order cognitive networks. More recently, Luppi et al.^33^ showed that cortical regions with higher HAR-BRAIN expression exhibit greater anesthesia-induced reductions in connectome identifiability, linking these genes to the neural basis of individual-specific functional connectivity. We examined whether regional HAR-BRAIN gene expression, quantified using transcriptional data from the Allen Human Brain Atlas (AHBA; human.brain-map.org), was associated with connectome fingerprinting. We hypothesized that cortical regions with higher HAR-BRAIN expression would exhibit stronger person-specific functional connectivity.

For each cortical parcel, HAR-BRAIN gene expression values were derived from the Allen Human Brain Atlas transcriptional data using the abagen Python package to map donor-level expression to the Schaefer 1,000-parcel atlas (Fig. 4A). Expression values were averaged across donors and summarized across the 415 HAR-BRAIN genes defined by Wei et al. (2019). Parcel-wise HAR-BRAIN expression was then correlated with fingerprinting accuracy using the same statistical procedures described above.

**Figure 4.**
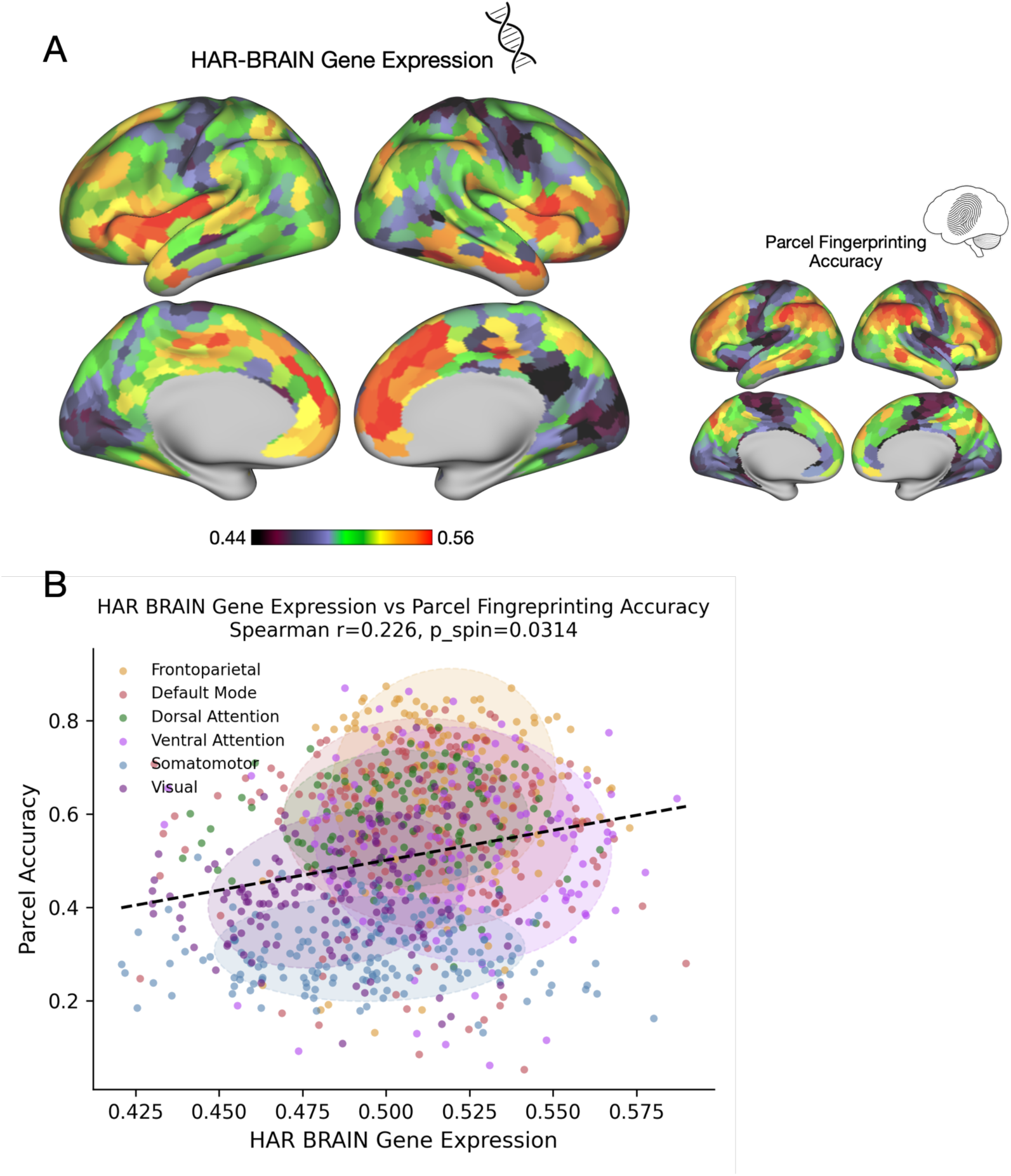
HAR-BRAIN gene expression is positively correlated with parcel-level fingerprinting accuracy. (A) Spatial distribution of HAR-BRAIN gene expression and parcel-wise fingerprinting accuracy mapped onto the Schaefer 1,000-parcel cortical surface. The fingerprinting accuracy map, as also shown in Fig. 2A, is reproduced at a smaller scale (right) for visual comparison. (B) Association between parcel-level HAR-BRAIN gene expression and fingerprinting accuracy. Each point represents one cortical parcel and is colored according to its canonical functional network. Shaded ellipses indicate the distribution of parcels within each canonical network, and dashed lines show the overall linear fit for visualization. Fingerprinting accuracy was positively and significantly associated with levels of HAR-BRAIN expression (*r* = 0.226, *p*_spin_ < 0.05).

As hypothesized, fingerprinting accuracy was positively associated with HAR-BRAIN gene expression (*r* = 0.226, *p_spin_* < 0.05; Fig. 4B). Cortical regions with greater expression of genes associated with human accelerated regions exhibited more individual-specific functional connectivity profiles. Analyses using Idiff yielded convergent results (Supplementary Figure S1).

## Functional homology with other primates

Across mammalian evolution, functional connectivity reflects both conserved network architecture and species-specific adaptations^34,35^. To quantify evolutionary conservation of human functional organization, Xu et al.^23^ developed the Functional Connectivity Homology Index (FCHI), a vertex-wise measure of cross-species similarity based on resting-state functional connectivity in humans and rhesus macaques^36^. FCHI is highest in primary visual and auditory cortices and lowest in higher-order association regions, reflecting a gradient of evolutionary divergence across the human cortex^23^. We therefore hypothesized that regions with lower functional homology, that is, with more human-specific functional organization, would exhibit greater fingerprinting accuracy than evolutionary conserved regions.

For each cortical parcel, mean functional homology was estimated by averaging the Xu et al.^23^ vertex-wise macaque–human FCHI values within the Schaefer 1,000-parcel atlas (Fig. 5A). Parcel-wise functional homology was then correlated with fingerprinting accuracy using the same statistical procedures described above.

**Figure 5.**
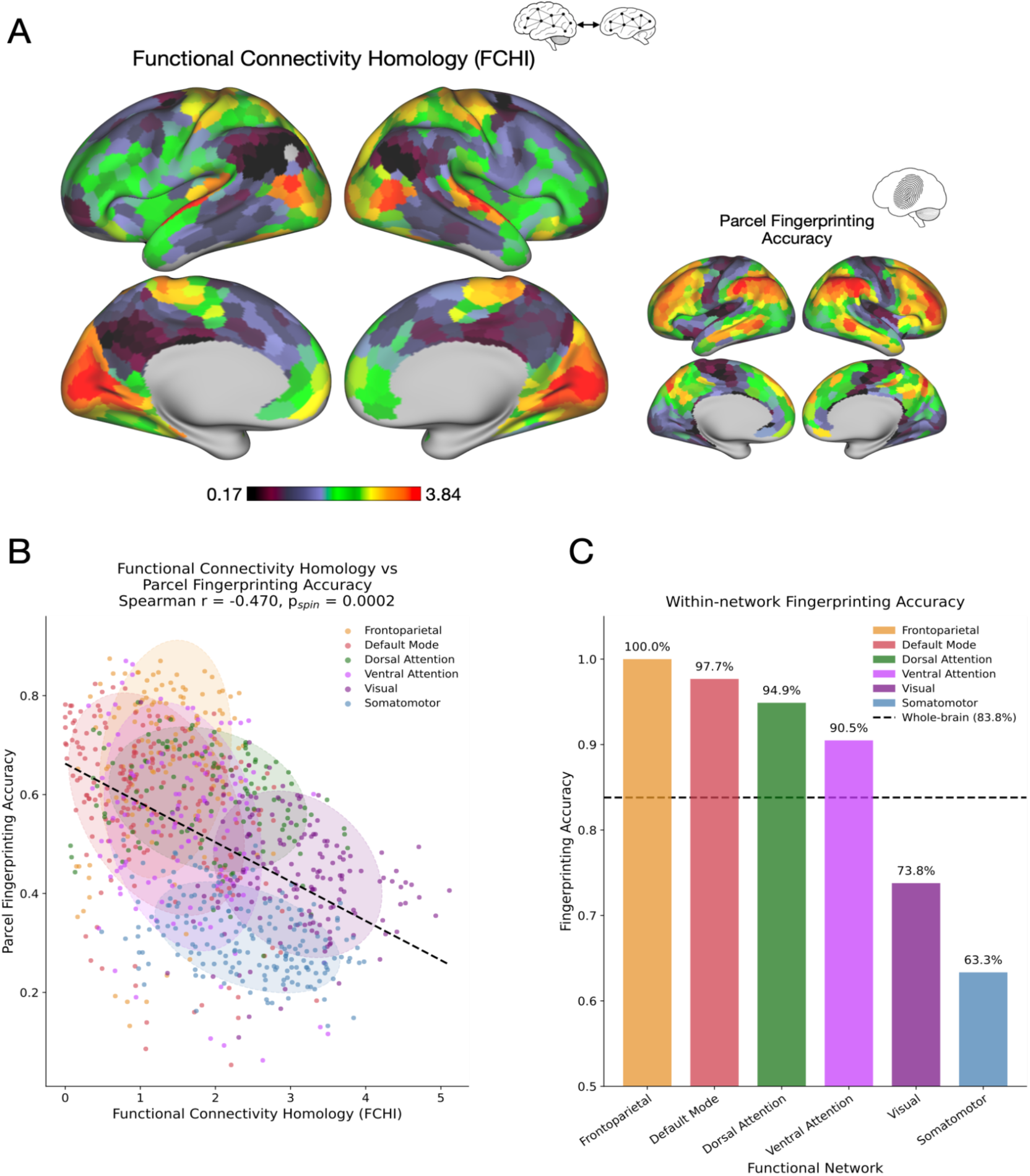
Functional homology is inversely associated with fingerprinting accuracy. (A) Spatial distribution of macaque-human functional homology and parcel-wise fingerprinting accuracy displayed on the Schaefer 1,000-parcel cortical surface. The fingerprinting-accuracy map, as also shown in Fig. 2A, is reproduced at a smaller scale for visual comparison with the functional homology map. Warmer colors indicate higher values. (B) Correlation between parcel-level functional homology and fingerprinting accuracy. Each point represents one cortical parcel and is colored according to its canonical functional network. Shaded ellipses indicate the distribution of parcels within each network, and the dashed line shows the overall linear fit for visualization. Fingerprinting accuracy was negatively and significantly associated with functional homology (*r* = −0.470, *p*_spin_ < 0.001). (C) Within-network fingerprinting accuracy across the six non-limbic canonical functional networks. Accuracy was highest in frontoparietal control and default mode networks and lowest in visual and somatomotor networks.

As hypothesized, fingerprinting accuracy was negatively associated with functional homology (*r* = −0.470, *p_spin_* < 0.001; Fig. 5B), indicating that cortical regions with less conserved functional organization across macaques and humans exhibited more distinctive individual-specific connectivity profiles. Analyses using Idiff yielded convergent results (Supplementary Figure S1).

To determine whether this association also emerged at the network level, we calculated fingerprinting accuracy separately within each of the canonical functional network using only intra-network functional edges. Fingerprinting accuracy was highest in the frontoparietal control network (100.0%), followed by the default mode (97.7%), dorsal attention (94.9%) and ventral attention (90.5%) networks, and was lower in the visual (73.8%) and somatomotor (63.3%) networks (Fig. 5C). This hierarchy closely mirrors the inverse gradient of cross-species functional homology reported by Xu et al.^23^. Association networks with lower macaque-human functional homology exhibited the highest fingerprinting accuracy, whereas evolutionary conserved sensory and motor networks exhibited the lowest. Because the number of within-network edges varied across networks, we repeated the analysis after matching the number of sampled edges. The network hierarchy remained unchanged, indicating that the observed differences were not attributable to network size or edge counts (Supplementary Fig. S2).

## Composite Evolutionary Index

The four complementary indices of human cortical evolutionary specialization described above are interrelated rather than independent to one another, we first quantified their pairwise spatial correlations (Figure 6A). Statistical significance was assessed using two-sided spatial spin tests and Benjamini–Hochberg false-discovery-rate correction across the six comparisons.

**Figure 6.**
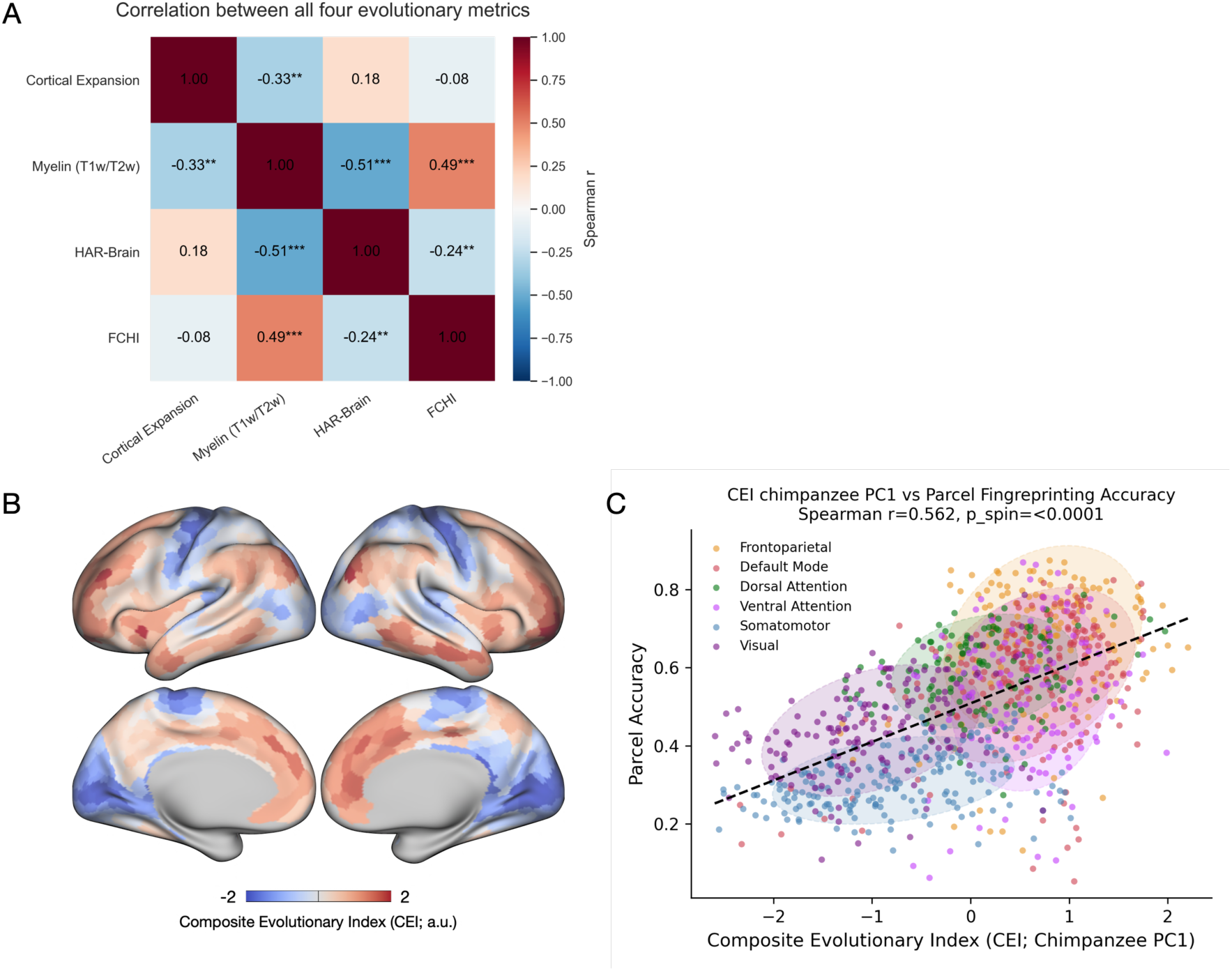
The Composite Evolutionary Index (CEI) and its association with parcel-level fingerprinting accuracy. (A) Pairwise correlations among chimpanzee-to-human cortical expansion, myelin (T1w/T2w), HAR-BRAIN gene expression and primate functional homology across cortical parcels. Asterisks indicate significance based on two-sided spatial spin tests followed by Benjamini– Hochberg false-discovery-rate correction for six comparisons: ∗ *p* < 0.05, ∗∗ *p* < 0.01 and ∗∗∗ *p* < 0.001. (B) Spatial distribution of the CEI and parcel-wise fingerprinting accuracy mapped onto the Schaefer 1,000-parcel cortical surface. (C) Association between parcel-wise CEI values and fingerprinting accuracy. Each point represents one cortical parcel and is colored according to its canonical functional network. Shaded ellipses indicate the distribution of parcels within each network, and the dashed line shows the overall linear fit for visualization. CEI was positively and significantly associated with fingerprinting accuracy (*r* = 0.562, *p_spin_* < 0.0001).

To capture shared spatial patterns across these evolutionary measures, we constructed a data-driven Composite Evolutionary Index (CEI) by integrating the four above-described metrics. We hypothesized that cortical regions with higher CEI values would exhibit greater fingerprinting accuracy. Parcel-wise values were standardized, and the directions of myelin and functional homology were reversed so that higher CEI values consistently reflect greater human-specific differentiation. Principal component analysis was then performed across cortical parcels, and the first principal component, which explained 49.1% of the variance across the four measures (using chimpanzee-to-human cortical expansion), was defined as the CEI. All four measures loaded positively on this component, with loadings of 0.517 for HAR-BRAIN expression, 0.337 for chimpanzee-to-human cortical expansion, 0.488 for reversed FCHI, and 0.617 for reversed myelin. The resulting CEI value for each cortical parcel is shown in Fig. 6B.

Parcel-wise CEI values were correlated with fingerprinting accuracy using a Spearman rank correlation, with statistical significance assessed using 10,000 spatial spin permutations to account for cortical spatial autocorrelation. As hypothesized, CEI was positively associated with fingerprinting accuracy (*r* = 0.562, *p_spin_* < 0.0001; Fig. 6C). Thus, cortical regions exhibiting more evolutionary specialized profile, characterized by greater chimpanzee-to-human expansion, higher HAR-BRAIN gene expression, lower myelin content (T1w/T2w), and lower functional homology, also exhibited more distinctive individual-specific functional connectivity profiles. Analyses using macaque-to-human cortical expansion to derive CEI, as well as analyses using Idiff yielded converging results (Supplementary Figure S3). Among all evolutionary measures examined, CEI showed the strongest association with fingerprinting accuracy.

## Discussion

Individuality is a fundamental feature of human biology and has long been recognized across philosophy, religious thought, psychology, and neuroscience^3,37–40^. From an evolutionary perspective, individual differences are expected to arise through the combined effects of genetic variation, development, epigenetic regulation and accumulated experience, yielding brains with distinctive structural and functional organization. Such variation provides the substrate upon which natural selection acts, increasing the likelihood that some individuals possess traits suited to changing environmental conditions^41^. Functional-connectome fingerprinting provides a neural expression of this principle: patterns of functional connectivity are sufficiently reproducible within individuals and variable between individuals to support person-level identification^3^.

Our findings extend this framework by showing that neural individuality is preferentially expressed in cortical territories that underwent pronounced modification during human evolution. Regions exhibiting the greatest expansion along the human lineage, or the strongest microstructural, molecular and functional divergence from other primates, were also those in which functional connectivity carried the most person-specific information. This spatial convergence raises the possibility that human cortical evolution increased not only cognitive capacity, but also the scope for individually differentiated functional architectures. Although these associations do not establish evolutionary causality, they link long-term cortical divergence across species to stable neural variation among living humans.

This relationship was strongest in higher-order association cortex. Fingerprinting accuracy peaked in the frontoparietal control network (FPCN) and default mode network (DMN), with the FPCN alone achieving 100% identification accuracy across all 431 participants. Connectivity within this network was therefore sufficiently reproducible within individuals and sufficiently distinctive between individuals to identify every member of the cohort. This result accords with previous studies showing that the FPCN and DMN contribute disproportionately to connectome fingerprinting^4^ and exhibit particularly high interindividual variability in functional organization^42^.

The prominence of these networks is also consistent with their roles in higher-order cognition. The DMN supports self-referential and internally generated cognition, whereas the FPCN flexibly coordinates distributed cortical systems during cognitive control, working memory, decision-making and goal-directed behavior^10,11^. That these systems are both evolutionarily differentiated and individually distinctive suggests that the cortical architecture supporting flexible and integrative cognition also provides a major substrate for neural individuality.

By contrast, primary visual, somatomotor and auditory cortices exhibited comparatively low fingerprinting accuracy, consistent with their more evolutionarily conserved organization. This distribution recapitulates the principal unimodal-to-transmodal gradient of cortical organization, which extends from primary sensory and motor systems to transmodal association regions, including the DMN^42^. Whereas unimodal cortices are specialized for domain-specific sensory and motor processing, transmodal cortices integrate information across systems to support higher-order cognition^9^. Individual specificity therefore appears to increase toward the transmodal end of the cortical hierarchy. Together with the correspondence to evolutionary maps, this pattern suggests that modifications of association cortex may have shaped not only capacities shared across humans, but also the dimensions along which individual brains differ.

Neural individuality, however, is not unique to humans. Individual-specific functional-connectivity signatures have been demonstrated in macaques^43^ and in phylogenetically distant species such as mice, in which both sensory and association networks support connectome-based identification^44^. Connectome individuality thus appears to be a broadly conserved feature of mammalian brain organization, although its anatomical expression may vary across species according to their genetic, developmental, ecological and experiential constraints. Extending the present comparative framework to species spanning greater phylogenetic distances, including mice and ferrets, could determine whether the correspondence between evolutionary divergence and individual specificity is strongest for recent human-lineage changes or reflects a deeper trajectory of mammalian cortical evolution.

Several limitations qualify the interpretation of these findings. First, the HAR-BRAIN gene set captures only one of several molecular signatures that distinguish the human cortex from that of other mammals. Zeng et al.^45^, for example, identified 19 human supragranular-enriched genes that are preferentially expressed in cortical layers II/III in humans but are absent from the mouse cortex or enriched primarily in deeper layers. Testing additional, independently defined molecular profiles will therefore be important for establishing the generality of the present associations. Second, the evolutionary measures examined here span different phylogenetic intervals. Whereas several indices capture human-specific differences or include comparisons with chimpanzees, the FCHI was derived from human–macaque comparisons and may reflect modifications accumulated over a substantially longer period. Convergence across these measures therefore cannot be attributed exclusively to the most recent stages of human evolution. Third, our analyses were restricted to the neocortex and did not include subcortical structures, the cerebellum or the brainstem. This analytic decision is based on previous literature showing little-to-no contribution of these regions to identification accuracy^3^. Whole-brain analyses optimized for these regions will be required to determine whether similar evolutionary principles extend beyond the neocortex.

In summary, our findings connect evolutionary variation across species with stable functional variation among individuals. Cortical regions that expanded or diverged most strongly in humans, particularly the frontoparietal control and default mode systems, also contained the connectivity patterns that most reliably distinguished one person from another, whereas more conserved unimodal regions were less individually distinctive. These results suggest that evolution did not generate neural individuality de novo, but may have amplified its expression within association cortex, where flexible integration, cognitive control, and self-related processing are concentrated. More broadly, the spatial correspondence between human cortical specialization and connectome uniqueness provides a framework for understanding how genetic, developmental, and experiential processes act on evolutionarily modified cortical systems to produce the distinctive functional architecture of each human brain.

## Materials and Methods

### Participants and imaging data

Resting-state functional MRI (rsfMRI) data were obtained from the publicly available Human Connectome Project Young Adult S1200 release (HCP)^5^. The present analysis included 431 participants with complete resting-state data from two imaging sessions conducted on separate days. Participants were randomly selected from each family unit such that no two participants are related. The two sessions, REST1 and REST2, were treated as repeated measurements for connectome fingerprinting, consistent with the test-retest framework introduced by Finn and colleagues^3^.

MRI data were acquired by the WU-Minn HCP using a customized 3 T Siemens Skyra scanner. The HCP resting-state protocol comprised approximately 30 min of rsfMRI per imaging session. Full acquisition parameters and quality-assurance procedures are described in the HCP protocol and associated publications^46^.

### Resting-state fMRI preprocessing

We used the rsfMRI data preprocessed and released by the HCP. Data underwent the HCP minimal preprocessing pipelines, including correction of gradient nonlinearity, head motion and EPI distortion; registration to each participant’s structural images; transformation to standard space; projection to the cortical surface; and representation in the CIFTI greyordinate framework^45^. The HCP ICA-FIX denoised rsfMRI data were used^47^.

### Cortical parcellation and functional connectivity matrices

The cerebral cortex was parcellated using the Schaefer 1,000-parcel atlas, with each parcel assigned to one of the seven canonical functional networks defined by Yeo and colleagues^17,18^. For each participant and session, the mean rsfMRI time series was extracted from each cortical parcel. Pairwise Pearson correlations were then calculated between all parcel time series and Fisher z-transformed, resulting in one 1,000 × 1,000 functional connectivity matrix per participant and session.

Only cortical parcels were included in all analyses. Parcels assigned to the Yeo limbic network were excluded to reduce potential bias caused by susceptibility-related signal loss and distortion in orbitofrontal and anterior temporal cortices^21,48^. The same parcel mask and ordering were used for all fingerprinting, evolutionary-map and spatial-null analyses.

### Parcel-wise connectome fingerprinting

Connectome fingerprinting was performed separately for each cortical parcel. For a given parcel, its row in the participant’s functional connectivity matrix (in other words, its functional connectivity with all other cortical parcels) was extracted as a parcel-specific connectivity profile. The parcel’s self-connection was excluded from this profile. Each participant was therefore represented by one connectivity profile for every parcel.

For each parcel, the REST1 connectivity profile of every participant was correlated with the corresponding parcel-specific REST2 profiles of all participants, producing a participant-by-participant cross-session similarity matrix. A participant was counted as successfully identified only when their within-participant correlation was the maximum value in both the corresponding row and column of the cross-session similarity matrix. Parcel-wise fingerprinting accuracy was defined as the proportion of participants correctly identified out of the whole sample.

Differential identifiability (Idiff) was calculated as a complementary continuous measure of fingerprinting strength. For each participant, the diagonal within-participant correlation was compared with the mean of the non-self correlations in the participant’s corresponding row and column. The participant-level difference was then averaged across participants to obtain a single Idiff value for each cortical parcel. Higher Idiff values indicate greater distinguishing power between self and others.

### Evolutionary and neurobiological cortical maps

All evolutionary and neurobiological measures were represented on the cortical surface and summarized within the Schaefer 1,000-parcel atlas. For each measure, vertex-wise values were averaged across all vertices belonging to each parcel.

### Macaque-to-human cortical expansion

The macaque-to-human cortical expansion map was obtained from Xu et al.^23^. In that work, cortical surface data from 48 rhesus macaques and 187 humans were used to establish cross-species correspondence, and local expansion was quantified as the ratio of human cortical surface area to the corresponding macaque surface area at each location on the human cortical surface.

### Chimpanzee-to-human cortical expansion

The chimpanzee-to-human cortical expansion map was obtained from Wei et al.^22^. Their analysis used pial-surface reconstructions from 29 chimpanzees and 50 humans to quantify regional cortical expansion.

### Cortical myelin

Throughout this study, the term myelin refers to T1w/T2w contrast, an MRI-based proxy for relative cortical myelin content rather than a direct histological measure^27^. We used the HCP group-average MSMAll-aligned cortical T1w/T2w myelin map.

### HAR-BRAIN gene expression

Regional HAR-BRAIN gene expression was estimated from post-mortem transcriptional data from the Allen Human Brain Atlas (AHBA)^49^. Tissue-sample data were processed and mapped to the Schaefer 1,000-parcel atlas using the abagen Python package^50^, with probes selected according to their differential stability across donors and donor-specific probe measurements aggregated before regional assignment. Sample-wise and gene-wise expression values were normalized using scaled robust sigmoid normalization, applied only to samples assigned to cortical, non-limbic parcels. Missing parcel values were estimated using spatial interpolation. Five AHBA donors were included (9861, 10021, 12876, 14380 and 15697); donor 15496 was excluded because the corresponding source data were unavailable. Expression values were then summarized across the 415 HAR-BRAIN genes defined by Wei et al.^22^ by z-scoring each gene across left-hemisphere non-limbic parcels and averaging the resulting scores, yielding one HAR-BRAIN expression value per parcel.

### Macaque-human functional homology

The functional homology map was obtained from Xu et al.^23^, who compared resting-state functional connectivity organization in rhesus macaques and humans and derived a regional Functional Connectivity Homology Index (FCHI). Higher FCHI values indicate greater similarity in functional network organization across species, whereas lower values indicate greater human-macaque differentiation.

### Associations between fingerprinting and cortical maps

Parcel-wise fingerprinting accuracy was related separately to macaque-to-human cortical expansion, chimpanzee-to-human cortical expansion, myelin, HAR-BRAIN gene expression and FCHI using Spearman rank correlations. Spearman correlations were selected because several cortical measures were non-normally distributed and because the hypotheses concerned monotonic spatial relationships.

Equivalent analyses using Idiff were conducted as supplementary analyses. The direction of each association was interpreted according to the biological meaning of the measure: positive associations were predicted for cortical expansion and HAR-BRAIN expression, whereas negative associations were predicted for myelin and FCHI. See Idiff association with all aforementioned metrics in Supplementary Figures S1.

### Spatial permutation testing

Statistical significance of parcel-wise spatial correlations was evaluated using 10,000 spherical spin permutations implemented via the alexander_bloch method in the neuromaps Python package^50^. Random hemisphere-preserving rotations were applied to the fsLR spherical surface, reassigning parcel values while preserving the spatial autocorrelation structure of cortical maps. For each permutation, the rotated fingerprinting map (Idiff or parcel accuracy) was correlated with the metric map. Two-sided spin-test P values were calculated as the proportion of null correlations whose absolute magnitude was at least as large as the observed correlation, with a +1 correction applied to both the numerator and denominator to avoid P = 0.

### Within-network fingerprinting

To assess whether cross-species functional homology was reflected at the level of canonical functional systems, we additionally calculated fingerprinting accuracy separately within each non-limbic Yeo network. For each network, participants were represented using only functional connections using pairs of parcels within that network. The same cross-session correlation and bidirectional identification procedure used in the parcel-wise analysis was then applied to each within-network connectivity profile.

### Edge-count quality control analysis

Because canonical networks differ in parcel number and therefore in the number of available within-network edges, we performed an edge-count-matched quality-control analysis. For each of seven target edge counts (250, 500, 1000, 2000, 4000, 6000, and 7,260), an equal number of within-network edges was randomly sampled from each non-limbic network, and fingerprinting accuracy was recalculated using the sampled edge sets (500 iterations per edge count; random seed = 42). Results are reported as the mean across iterations with standard error of the mean ribbons. The relative ordering of networks remained stable across sampled edge counts, indicating that the observed network hierarchy was not driven by differences in network size or edge number. See Supplementary Figure S2.

### Composite Evolutionary Index

To capture the shared spatial pattern of the four evolutionary and neurobiological measures, we constructed a Composite Evolutionary Index (CEI). The primary CEI integrated chimpanzee-to-human cortical expansion, myelin, HAR-BRAIN gene expression and macaque-human functional homology.

Each parcel-wise measure was standardized across retained parcels. The signs of myelin and FCHI were flipped so that higher values consistently indicated greater evolutionary differentiation in humans: greater cortical expansion, higher HAR-BRAIN expression, lower myelin and lower cross-species functional homology. Principal component analysis was then applied to the four standardized measures across parcels. Scores on the first principal component were used as the CEI and reoriented, when necessary, so that higher scores corresponded to the evolutionarily derived direction defined above. CEI scores were subsequently standardized across parcels. For the macaque-based CEI, the first principal component explained 51.0% of the variance across the four standardized measures. After orienting the component so that higher scores reflected greater evolutionary differentiation, the component loadings were 0.431 for HAR-BRAIN expression, 0.416 for macaque-to-human cortical expansion, 0.584 for reversed FCHI and 0.548 for reversed myelin. For the chimpanzee-based CEI, the first principal component explained 49.1% of the variance, with corresponding loadings of 0.517, 0.337, 0.488 and 0.617, respectively.

The relationship between the chimpanzee-based CEI and parcel-wise fingerprinting accuracy was assessed using a Spearman rank correlation and 10,000 spatial spin permutations. A complementary CEI replacing chimpanzee-to-human expansion with macaque-to-human expansion was evaluated in the Supplementary Information (See Supplementary Figure S3). Equivalent CEI analyses using Idiff were also conducted as supplementary analyses (Supplementary Figure S3).

#### Software and reproducibility

Analyses were performed in Python using NumPy, pandas, SciPy, nibabel, h5py, matplotlib and, where available, netneurotools. AHBA processing was performed using abagen. CIFTI surface files were handled in fsLR space.

## Data availability

All data analysed in this study are publicly available from existing resources. Resting-state functional MRI data were obtained from the Human Connectome Project Young Adult S1200 release, and post-mortem gene-expression data were obtained from the Allen Human Brain Atlas. No new primary data were generated in this study.

## Code availability

Custom MATLAB and Python code used to generate the functional connectivity matrices, perform connectome fingerprinting analyses, construct the Composite Evolutionary Index, conduct spatial permutation tests and generate the reported figures is publicly available on GitHub at https://github.com/nogayair/Evolutionary_Fingerprinting.

## Acknowledgements

Funding statements

This research was supported by the European Research Council (ERC-2023-ADG 101141436).

## Author contribution statements

Conceptualization: NY, YC, YBH, IT

Methodology: NY, YC

Investigation: NY, YC

Visualization: NY, YC

Supervision: YBH, IT

Writing - original draft, review: NY, YC

Writing - editing: NY, YC, YBH, IT

## References

1. Andrews-Hanna, J. R., Smallwood, J. & Spreng, R. N. The default network and self-generated thought: component processes, dynamic control, and clinical relevance. Ann. N. Y. Acad. Sci. 1316, 29–52 (2014).

2. Di Plinio, S., Perrucci, M. G., Aleman, A. & Ebisch, S. J. H. I am Me: Brain systems integrate and segregate to establish a multidimensional sense of self. NeuroImage 205, 116284 (2020).

3. Finn, E. S. et al. Functional connectome fingerprinting: identifying individuals using patterns of brain connectivity. Nat. Neurosci. 2015 1811 18, 1664–1671 (2015).

4. Amico, E. & Goñi, J. The quest for identifiability in human functional connectomes. Sci. Rep. 2018 81 8, 1–14 (2018).

5. Van Essen, D. C. et al. The WU-Minn Human Connectome Project: An overview. NeuroImage 80, 62–79 (2013).

6. Sorrentino, P. et al. Clinical connectome fingerprints of cognitive decline. NeuroImage 238, 118253 (2021).

7. Stampacchia, S. et al. Fingerprints of brain disease: connectome identifiability in Alzheimer’s disease. *Commun*. Biol. 7, 1169 (2024).

8. Yair, N. et al. Experience-dependent changes in functional connectome fingerprinting. Nat. Commun. 10.1038/s41467-026-77094-y (2026) doi:10.1038/s41467-026-77094-y.

9. Kringelbach, M. L. & Deco, G. Prefrontal cortex drives the flexibility of whole-brain orchestration of cognition. Curr. Opin. Behav. Sci. 57, 101394 (2024).

10. Seitzman, B. A. et al. Trait-like variants in human functional brain networks. Proc. Natl. Acad. Sci. 116, 22851–22861 (2019).

11. Duncan, J. The multiple-demand (MD) system of the primate brain: mental programs for intelligent behaviour. Trends Cogn. Sci. 14, 172–179 (2010).

12. Cole, M. W. et al. Multi-task connectivity reveals flexible hubs for adaptive task control. Nat. Neurosci. 16, 1348–1355 (2013).

13. Buckner, R. L. & Krienen, F. M. The evolution of distributed association networks in the human brain. Trends Cogn. Sci. 17, 648–665 (2013).

14. Hill, J. et al. Similar patterns of cortical expansion during human development and evolution. Proc. Natl. Acad. Sci. 107, 13135–13140 (2010).

15. Zilles, K., Armstrong, E., Schleicher, A. & Kretschmann, H.-J. The human pattern of gyrification in the cerebral cortex. Anat. Embryol. (Berl.) 179, 173–179 (1988).

16. Gibbs, R. A. et al. Evolutionary and Biomedical Insights from the Rhesus Macaque Genome. Science 316, 222–234 (2007).

17. Schaefer, A. et al. Local-Global Parcellation of the Human Cerebral Cortex from Intrinsic Functional Connectivity MRI. Cereb. Cortex 28, 3095–3114 (2018).

18. Thomas Yeo, B. T., et al. The organization of the human cerebral cortex estimated by intrinsic functional connectivity. J. Neurophysiol. 106, 1125–1165 (2011).

19. Barton, R. A. & Harvey, P. H. Mosaic evolution of brain structure in mammals. Nature 405, 1055–1058 (2000).

20. Rakic, P. Evolution of the neocortex: a perspective from developmental biology. Nat. Rev. Neurosci. 10, 724–735 (2009).

21. Girn, M., Setton, R., Turner, G. R. & Spreng, R. N. The “limbic network,” comprising orbitofrontal and anterior temporal cortex, is part of an extended default network: Evidence from multi-echo fMRI. Netw. Neurosci. 8, 860–882 (2024).

22. Wei, Y. et al. Genetic mapping and evolutionary analysis of human-expanded cognitive networks. Nat. Commun. 10, 4839 (2019).

23. Xu, T. et al. Cross-species functional alignment reveals evolutionary hierarchy within the connectome. NeuroImage 223, 117346 (2020).

24. Glasser, M. F., Goyal, M. S., Preuss, T. M., Raichle, M. E. & Van Essen, D. C. Trends and properties of human cerebral cortex: Correlations with cortical myelin content. NeuroImage 93, 165–175 (2014).

25. Essen, D. C. V. & Dierker, D. L. Surface-Based and Probabilistic Atlases of Primate Cerebral Cortex. Neuron 56, 209–225 (2007).

26. Donahue, C. J., Glasser, M. F., Preuss, T. M., Rilling, J. K. & Van Essen, D. C. Quantitative assessment of prefrontal cortex in humans relative to nonhuman primates. Proc. Natl. Acad. Sci. 115, E5183–E5192 (2018).

27. Glasser, M. F. & Essen, D. C. V. Mapping Human Cortical Areas In Vivo Based on Myelin Content as Revealed by T1- and T2-Weighted MRI. J. Neurosci. 31, 11597–11616 (2011).

28. Florio, M. et al. Human-specific gene ARHGAP11B promotes basal progenitor amplification and neocortex expansion. Science 347, 1465–1470 (2015).

29. Oldham, M. C., Horvath, S. & Geschwind, D. H. Conservation and evolution of gene coexpression networks in human and chimpanzee brains. Proc. Natl. Acad. Sci. 103, 17973–17978 (2006).

30. Cáceres, M. et al. Elevated gene expression levels distinguish human from non-human primate brains. Proc. Natl. Acad. Sci. 100, 13030–13035 (2003).

31. O’Leary, D. D. M., Chou, S.-J. & Sahara, S. Area Patterning of the Mammalian Cortex. Neuron 56, 252–269 (2007).

32. Pollard, K. S. et al. An RNA gene expressed during cortical development evolved rapidly in humans. Nature 443, 167–172 (2006).

33. Luppi, A. I. et al. General anaesthesia decreases the uniqueness of brain functional connectivity across individuals and species. *Nat*. Hum. Behav. 9, 987–1004 (2025).

34. Griffa, A. et al. Evidence for increased parallel information transmission in human brain networks compared to macaques and male mice. Nat. Commun. 14, 8216 (2023).

35. Mantini, D., Corbetta, M., Romani, G. L., Orban, G. A. & Vanduffel, W. Evolutionarily Novel Functional Networks in the Human Brain? J. Neurosci. 33, 3259–3275 (2013).

36. Milham, M. P. et al. An Open Resource for Non-human Primate Imaging. Neuron 100, 61–74.e2 (2018).

37. Mishnah Sanhedrin 4:5. https://www.sefaria.org.il/Mishnah_Sanhedrin.4.5.

38. Allport, G. W. Pattern and Growth in Personality. xiv, 593 (Holt, Reinhart & Winston, Oxford, England, 1961).

39. Olson, M. V. Human Genetic Individuality. Annu. Rev. Genomics Hum. Genet. 13, 1–27 (2012).

40. Leibniz, G. W. Discourse on Metaphysics. in Philosophical Papers and Letters (eds Leibniz, G. W. & Loemker, L. E.) 303–330 (Springer Netherlands, Dordrecht, 1989). doi:10.1007/978-94-010-1426-7_36.

41. Barrett, R. D. H. & Schluter, D. Adaptation from standing genetic variation. Trends Ecol. Evol. 23, 38–44 (2008).

42. Margulies, D. S. et al. Situating the default-mode network along a principal gradient of macroscale cortical organization. Proc. Natl. Acad. Sci. 113, 12574–12579 (2016).

43. Miranda-Dominguez, O. et al. Connectotyping: Model Based Fingerprinting of the Functional Connectome. PLOS ONE 9, e111048 (2014).

44. Bergmann, E., Gofman, X., Kavushansky, A. & Kahn, I. Individual variability in functional connectivity architecture of the mouse brain. *Commun*. Biol. 3, 738 (2020).

45. Zeng, H. et al. Large-Scale Cellular-Resolution Gene Profiling in Human Neocortex Reveals Species-Specific Molecular Signatures. Cell 149, 483–496 (2012).

46. Glasser, M. M. F. et al. The minimal preprocessing pipelines for the Human Connectome Project. NeuroImage 80, 105–124 (2013).

47. Salimi-Khorshidi, G. et al. Automatic denoising of functional MRI data: Combining independent component analysis and hierarchical fusion of classifiers. NeuroImage 90, 449–468 (2014).

48. Weiskopf, N., Hutton, C., Josephs, O., Turner, R. & Deichmann, R. Optimized EPI for fMRI studies of the orbitofrontal cortex: compensation of susceptibility-induced gradients in the readout direction. Magn. Reson. Mater. Phys. Biol. Med. 20, 39–49 (2007).

49. Hawrylycz, M. J. et al. An anatomically comprehensive atlas of the adult human brain transcriptome. Nature 489, 391–399 (2012).

50. Markello, R. D. et al. Standardizing workflows in imaging transcriptomics with the abagen toolbox. eLife 10, e72129 (2021).

