## Supplemental Figures for "Evolution and Human Neural Individuality"

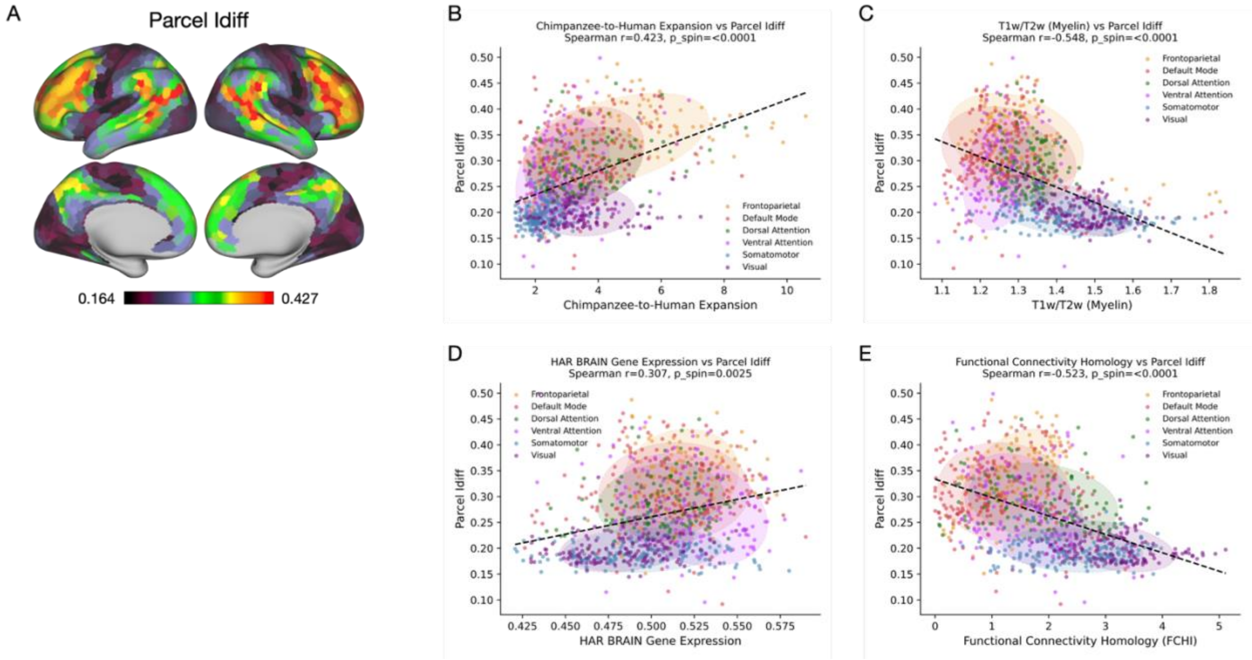


**Supplementary Figure S1. Reproduction of results with Idiff.** (A) Parcel-wise fingerprinting Idiff scores. Scatter plots presents associations between fingerprinting Idiff and parcel-level (B) chimpanzee-to-human cortical expansion; (C) myelin content (T1w/T2w); (D) HAR-BRAIN gene expression levels; and functional connectivity homology with macaques. Each point represents one cortical parcel and is colored according to its canonical functional network. Shaded ellipses indicate the distribution of parcels within each canonical network, and dashed lines show the overall linear fit for visualization. All four evolutionary measures were significantly associated with Idiff.

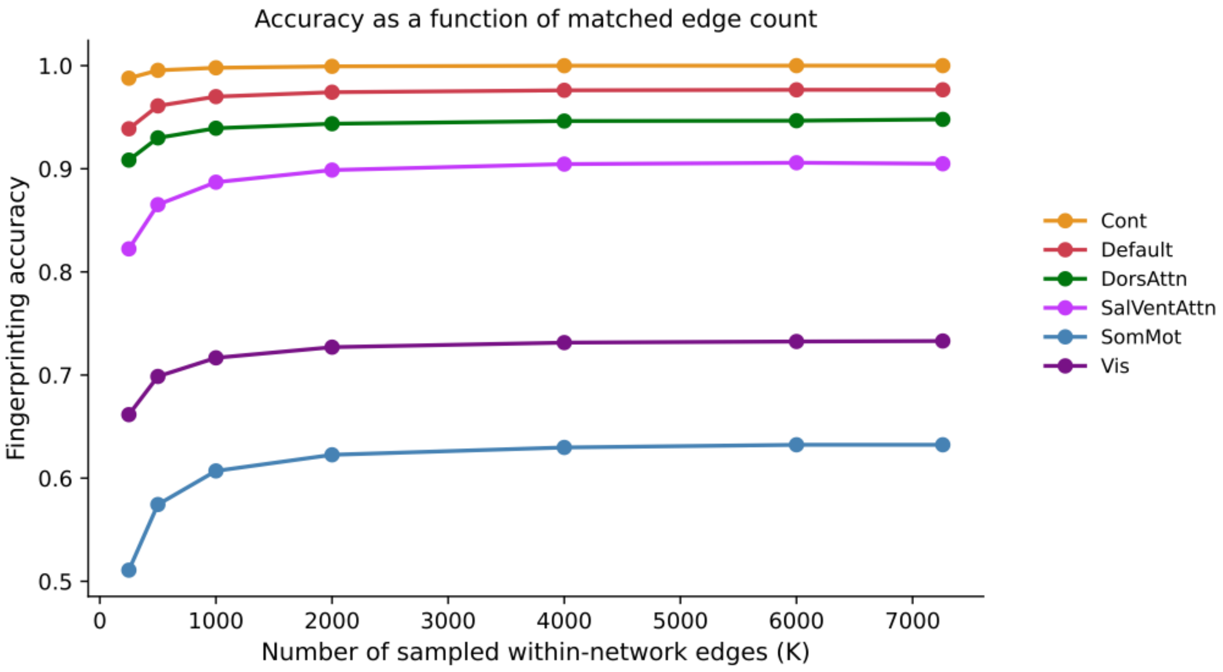


**Supplementary Figure S2. Fingerprinting accuracy with matched number of edges across networks.** Within-network fingerprinting accuracy across the six non-limbic canonical functional networks, with 250 (left) to 7260 (right) edges randomly selected withinp each network. Order of networks remained consistent across number of edges, with the highest accuracy exhibited in the frontoparietal control and default mode networks and lowest in visual and somatomotor networks.

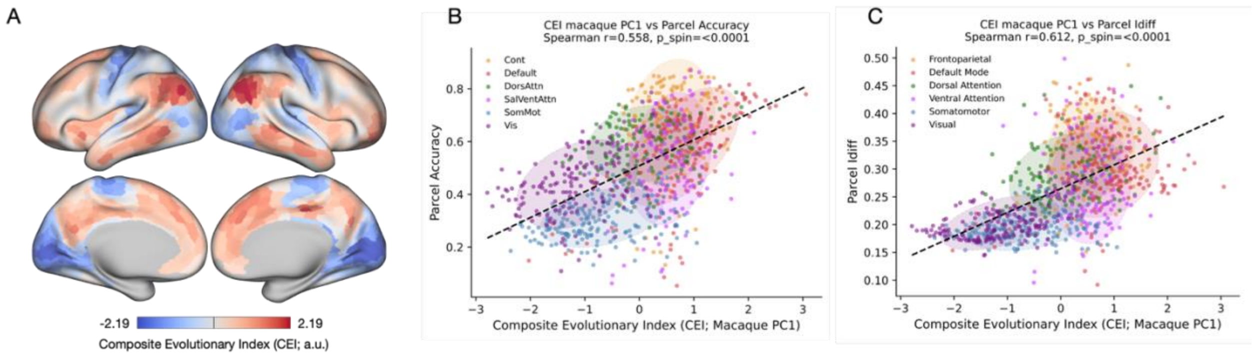


**Supplementary Figure S3. The Composite Evolutionary Index (CEI) using macaque-to-human cortical expansion** (A) Spatial distribution of the CEI and parcel-wise fingerprinting accuracy mapped onto the Schaefer 1,000-parcel cortical surface. (B) Association between parcel-wise CEI values and fingerprinting accuracy. (C) Association between parcel-wise CEI values and fingerprinting Idiff. Each point represents one cortical parcel and is colored according to its canonical functional network. Shaded ellipses indicate the distribution of parcels within each network, and the dashed line shows the overall linear fit for visualization. CEI was positively and significantly associated with fingerprinting accuracy ($r = 0.558$, $p_{spin} < 0.0001$) and Idiff ($r = 0.612$, $p_{spin} < 0.0001$).
